# Is that clear? Robust electrophysiological measures of the effects of prior knowledge on degraded speech perception

**DOI:** 10.64898/2026.07.31.742060

**Authors:** Shyanthony R. Synigal, Wen Li, Megan R. Serody, Judy L. Thompson, Edmund C. Lalor

## Abstract

Perception and sensation are not synonymous. Rather, perception is a process whereby sensory input is organized and interpreted in a behaviorally relevant way based on memory, experience, and context. One specific framework that is commonly invoked to explain perception is that of Bayesian inference. This framework casts perception as a probabilistic process whereby imprecise sensory data are combined with prior knowledge (prior) to determine what is consciously perceived (the posterior probability), which reflects the brain’s best guess as to the cause(s) of the sensory data. A striking behavioral example of how prior information can influence perception is seen in studies in which degraded speech is rendered intelligible by presenting information about the speech content in advance. Neurophysiological studies of this phenomenon have primarily focused on how it affects neural indices of low-level sensory encoding. The size of any reported effects on these indices tends to be much smaller – and much less consistent – than the notably large effects on perception that come with prior information. In the present study, we recorded EEG from 27 healthy adult participants (16 female) as they listened to degraded speech clips that were preceded by matching or mismatching text. Prior knowledge in the form of matching text led to a large perceptual pop-out effect when listening to degraded speech. Analyses of the resulting EEG revealed: 1) significant but relatively weak effects of prior information on EEG measures of the linguistic encoding of speech; and 2) a very large effect of prior information on an EEG signal that resembles a well-established neural index of perceptual evidence accumulation and that was strongly related to speech intelligibility ratings across participants. These EEG signals likely relate to separate components of a Bayesian inferential process during the predictive perception of degraded speech. As such, they have implications for understanding predictive perception more broadly and for future research on perceptual disturbances in clinical populations.

## INTRODUCTION

It is increasingly accepted that perception is the process of inferring the causes of our sensory input by combining that input with prior knowledge (Clark, 2013; Hohwy, 2013). This prior knowledge can come in the form of long-term knowledge that is “hard-wired” into our brains by evolution and learning (Knill & Pouget, 2004) or from the immediate context, wherein recent information can profoundly influence what we perceive when presented with new sensory input (Davis & Johnsrude, 2007).

A wealth of research over the years has suggested that the influence of prior information on perception can be accounted for by a process of Bayesian inference (Knill & Pouget, 2004). Under this account, perception is an active inferential process wherein the brain settles on its best guess as to the causes of its sensory input by combining that sensory input with prior knowledge. In the language of Bayesian inference, the noisy sensory input (or, rather, the noisy neural representation of that sensory input) contributes to the likelihood function, which gets combined with prior information (or predictions) to produce a percept that represents the posterior probability of the cause of the sensory input (Friston, 2005; Knill & Pouget, 2004; Rao & Ballard, 1999).

One powerful example of perceptual inference in action is the pop-out phenomenon that people experience when they are presented with heavily degraded speech that is preceded by informative prior information (Davis & Johnsrude, 2007; Remez et al., 1981; Sohoglu et al., 2012). Across many studies, researchers have shown that when listeners are provided with knowledge about the content of upcoming degraded speech, they report that they can hear that speech much more clearly. This effect has been shown when prior knowledge is provided as a clear audio version of the upcoming degraded speech (Di Liberto, Crosse, et al., 2018; Karunathilake et al., 2023), as a written text version of the upcoming degraded speech (Corcoran et al., 2023; Sohoglu & Davis, 2020; Sohoglu et al., 2012), or even semantically coherent preceding context (Signoret et al., 2018).

While the effect size of this behavioral pop-out phenomenon has been shown to be large and reliable across participants (Sohoglu et al., 2014), electrophysiological markers of the effect have generally been quite weak and have been rather inconsistent across studies. For example, some EEG/MEG indices of acoustic speech processing (e.g., cortical tracking of the speech envelope or spectrogram) have shown stronger responses to the acoustic features of degraded speech when it has been rendered intelligible by informative priors (Baltzell et al., 2017; Corcoran et al., 2023; Di Liberto, Lalor, et al., 2018; Holdgraf et al., 2016; Peelle et al., 2013). However, others have failed to find such an effect on similar measures of low-level speech processing, despite large behavioral pop-out (Di Liberto, Crosse, et al., 2018; Karunathilake et al., 2023; Millman et al., 2015). Similarly, neural indices reflecting the categorization of speech sounds into discrete phonetic classes have shown differences between informative and noninformative priors in some studies (Di Liberto, Crosse, et al., 2018), but not others (Karunathilake et al., 2023).

One possible explanation for the relatively weak and inconsistent electrophysiological measures of speech encoding in the literature is that the behavioral pop-out phenomenon likely relates more closely to the posterior of an inferential process (i.e., the percept), while the majority of electrophysiological studies of the phenomenon have focused on how neural data encodes the stimulus. Such neural measures are likely to relate much more closely to the likelihood function of a Bayesian process and may be only relatively weakly affected by prior information via hierarchical predictions as proposed by the predictive coding theory (Friston & Kiebel, 2009). Some evidence in support of this was provided recently in studies showing that neural markers of word level processing are significantly enhanced for degraded speech that has been made intelligible with valid priors (Karunathilake et al., 2023; Synigal et al., 2026). Responses to word onsets in continuous speech are not fully explainable based on the acoustics of that speech and, therefore, must surely – at least partially – reflect the cognitive process of perceiving words as words. As such, these effects, which were large for some specific measures (d = 0.89); (Karunathilake et al., 2023) are likely much more closely linked to perception and the Bayesian posterior than many of those mentioned above.

The goals of the present study are to examine neural signatures of sensory encoding and to search for neurophysiological evidence that is more directly related to the process of perceptual inference. To do this, we cast the problem as one of perceptual evidence accumulation based on uncertain sensory inputs. Specifically, we derive EEG measures of sensory and linguistic processing and search for EEG measures of perceptual evidence accumulation based on the read out of that sensory and linguistic processing. We report neurophysiological indices of such evidence accumulation that show unprecedentedly large effect sizes that mirror those seen in behavior.

## METHODS

### Participants

Data from 29 healthy adults (16 females, mean age = 22.759, SD = 4.094) were collected in this study. One participant was excluded due to an insufficient amount of data, and a second was excluded because they were unable to perform the task, resulting in a data set of 27 participants. Each participant reported having normal hearing, normal or corrected-to-normal vision, English as their first and main language, and no history of neurological disorders, claustrophobia, or hyperacusis. Participants provided written informed consent beforehand and were compensated for their participation. All procedures were approved by the University of Rochester Human Subjects Review Board.

### Stimuli and experimental procedure

The participants completed a perceptual pop-out experiment involving degraded speech. Specifically, they listened to three-channel noise-vocoded sentences that were preceded by the presentation of a matching or mismatching sentence as text on a screen. Their task was to read the text of the preceding sentence, listen to the degraded audio sentence, and then rate their ability to hear and understand words in the audio (**Figure 1**). A text prior was used here to mitigate any effects that might be caused by acoustic repetition (Di Liberto, Crosse, et al., 2018; Grill-Spector et al., 2006; Todorovic et al., 2011). The sentences were 2.35 s (±0.29 s) long on average and participants were provided with an amount of time to read each preceding sentence that was equal to the duration of the audio sentence plus an additional two seconds (i.e., 4.35 s on average). Participants were asked to rate the clarity of each audio sentence immediately following its presentation. Specifically, they were asked to rate the clarity based on a scale of 1-5, where 1 indicated that they did not detect any words in the audio; 2 indicated that they thought it sounded like there were words in the audio, but couldn’t understand them; 3 indicated they understood one or two words; 4 indicated they understood three or more words; and 5 indicated they understood a complete sentence. Each participant first performed a practice session which consisted of 12 trials and were given the opportunity to repeat the practice trials if needed.

**Figure 1.**
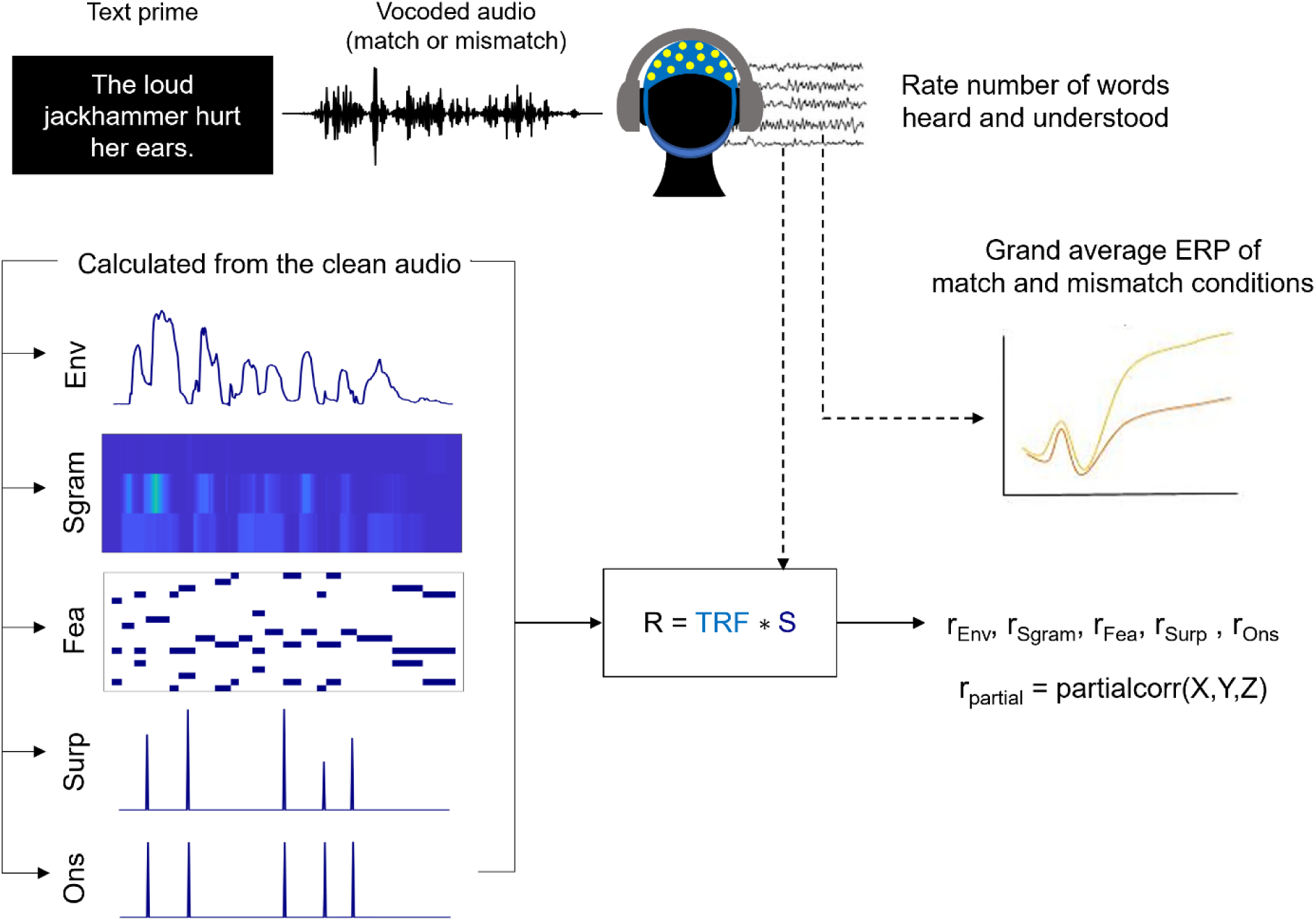
Methods. EEG data were recorded while participants listened to noise-vocoded speech (2.35 sec on average) preceded by matching or mismatching text. Afterwards, they rated their ability to hear and understand words in the clip. Forward modeling was used to estimate EEG responses (R) from the speech envelope, spectrogram, phonetic features, and word surprisal (S) features. Model performance (r) was assessed by calculating the correlation between the predicted EEG and the actual EEG data. Grand-average event-related potentials were also computed separately for matched and mismatched prior conditions. This was done by averaging the EEG (including low frequencies) across trials within each condition time-locked to the onset of the degraded speech.

Our instructions to participants were quite specific. Even though our task had essentially only two conditions (text and audio match or text and audio mismatch), we did not want participants to judge “match” vs “mismatch” as this would not necessarily inform us as to the clarity with which they were perceiving the speech. This is why we specifically instructed them to focus on the number of words they were able to hear and understand in the audio (regardless of whether the text and audio matched). In the main experiment, each participant completed blocks of 20 trials, with 10 match trials and 10 mismatch trials in a randomized order. An optional break was given after each block, and a mandatory break was taken after each quarter of the experiment (based on the number of possible trials) was completed. There were 500 possible trials for 6 participants, and 400 possible trials for the remaining participants. The EEG session time was limited, so participants completed as many trials as they could before the session ended (ranging from 160-320 trials).

A mixture of sentence sources was used for this experiment: a subset of Harvard sentences (Kabal, 2002), Bamford-Kowal-Bench (BKB) sentences (Bench et al., 1979) modified by the authors, and additional sentences created by the authors. A few of the Harvard sentences contained large volume fluctuations, so we applied MATLAB’s dynamic range compressor with a -27dB threshold, 7dB knee width, and a compression ratio of 15 to equalize the volume in those clips. The BKB sentences were modified to extend their time length and provide more semantic content. The experimenters also created sentences from scratch and attempted to match the length of the Harvard and modified BKB sentences. We recorded one of the authors reciting the modified and self-created sentences at 48kHz using the Blue Yeti USB microphone (https://www.logitech.com/) and Audacity (version 3.1.3) in a soundproof booth. Due to a technical error, a subset of sentences was recorded at 44.1kHz.

Three-channel noise vocoding was applied to each sentence. To do so, we first filtered the clean speech and white noise into three logarithmically spaced frequency bands according to Greenwood’s equation (70-602-2338-8000 Hz). We extracted the envelope of the three clean bands by calculating the absolute value of their Hilbert transforms. We then “compressed” the envelopes by computing the square root of their values. This was done to make the resulting stimuli more difficult to understand (Wild et al., 2012), in the hope of avoiding ceiling effects. Next, the envelopes were low pass filtered at 30 Hz using a 2^nd^ order Butterworth filter. We then modulated the filtered noise by the envelopes in the corresponding frequency bands and summed the resulting bands together. The stimuli were presented through Sennheiser HD650 headphones using Psychtoolbox (Kleiner et al., 2007) and custom MATLAB scripts. Both experiments took place in a dark soundproof booth.

### Data acquisition and preprocessing

We acquired 128 channel EEG (plus two mastoid channels) at a 512 Hz sampling rate with the BioSemi Active Two system. The data were preprocessed in two different ways: one to prepare the data for analyses aimed at identifying neural signatures of acoustic and linguistic speech processing, and a second to prepare the data for analysis aimed at identifying evidence of perceptual evidence accumulation.

The first preprocessing pipeline—aimed at preparing to model how the EEG responses to speech related to different acoustic and linguistic features of that speech—involved first attenuating slow drifts in the data by detrending the EEG at 1 Hz using the PREP pipeline *removeTrend* function (Bigdely-Shamlo et al., 2015). Noisy channels were then identified based on kurtosis, probability, and the mean standard deviation of the 8 neighboring channels (thresholds of 5, 5, and 3, respectively) and were recalculated via spherical spline interpolation (Delorme & Makeig, 2004). To further reduce artifacts, artifact subspace reconstruction (Kothe & Makeig, 2013) was applied using the EEGLAB *clean_rawdata* plugin with a standard deviation threshold of 20. The data were subsequently re-referenced to the average of the mastoid channels, and independent component analysis was performed using EEGLAB’s *picard* function to remove eye, muscle, line noise, and channel artifacts (0.8, 0.8, 0.9, 0.9 thresholds, respectively). Finally, the cleaned data were lowpass filtered at 8 Hz using a finite impulse response Kaiser window (10 Hz cutoff frequency, 1 dB passband attenuation, 60 dB stopband attenuation), epoched, and downsampled to 128 Hz.

The second preprocessing pipeline was aimed at identifying (likely slow) neural signatures of perceptual evidence accumulation and was done using custom MATLAB scripts applied to the original raw data at 512 Hz. First, all 128 scalp EEG channel were re-referenced to the average of the two mastoid channels prior to further processing. The re-referenced signals were low-pass filtered at 40 Hz using a zero-phase, fourth-order Butterworth filter. Bad channels were determined by visual inspection and interpolated using spherical interpolation as implemented in EEGLAB (Delorme & Makeig, 2004).

### Modeling the relationship between speech features and EEG responses

We sought to model how the EEG responses to the degraded speech stimuli related to different features of those speech stimuli. We focused on specific speech features that have been used in previous vocoding studies – under the assumption that different features may be differentially affected by prior information. Specifically, we selected a range of hierarchical speech features, each of which were calculated on the clean versions of the audio clips.

#### Envelope and spectrogram

The clean speech stimuli were lowpass filtered at 20 kHz (22.05 kHz cutoff frequency, 1 dB passband attenuation, 60 dB stopband attenuation). A gammachirp auditory filterbank was used to calculate the broadband envelope. This filterbank was used to filter the speech into a 3-band spectrogram from 70 Hz to 8 kHz with an equal loudness contour (Irino & Patterson, 2006). The frequency bands were averaged together, resulting in the speech envelope.

#### Phonetic features

The Montreal Forced Aligner (McAuliffe et al., 2017) was used to partition and time align each word in the story into phonemes according to the International Phonetic Alphabet (IPA) for American English. This software finds the onset and offset times for each word and phoneme. We then linearly mapped each phoneme onto a set of 19 phonetic features relating to manner of articulation, place of articulation, voicing of a consonant, backness of a vowel, and diphthongs.

#### Lexical surprisal

Lexical surprisal was calculated for each word using the Transformer-XL language model. This model is trained on a text corpus to predict the next word probability of an upcoming word using the context from all preceding words. We calculated the negative log of each word’s probability to estimate word surprisal (Dai et al., 2019). This feature was represented as a series of impulses, with a single impulse at each word onset whose height corresponded to the lexical surprisal value. To dissociate surprisal from word onset responses, a separate word onset regressor was constructed, consisting of an impulse (amplitude = 1) located at each word onset.

We wished to assess how the encoding of these different hierarchical speech features was affected by prior information during degraded speech processing. To that end, we used forward modeling to predict participants’ neural responses from the speech envelope, spectrogram, phonetic features, lexical surprisal, and word onset features. The EEG and each speech feature were z-scored prior to modeling. For each condition (i.e., separately for the match and mismatch trials), five-fold nested cross-validation was used to select the optimal ridge parameter (10^-3^-10^5^). For each fold, one partition was held out as test data, and the remaining data were used for training. Within the training set, cross-validation (via *mTRFcrossval*) was used to identify the ridge parameter that maximized prediction accuracy. The forward models, or temporal response functions (TRFs), were then computed using the selected ridge parameter and the training data. The neural response, r(t, n), sampled at times *t = 1…T* with *n* electrodes, is the result of channel specific TRFs, w(τ, n), convolved with the lagged stimulus features, s(t − τ), in addition to residual responses not explained by the model, ε(t, n). The lags ranged from 0 to 650 ms to capture early and late latency neural responses. This stimulus-response mapping is represented as (Crosse et al., 2016):

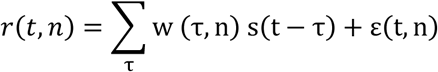

The goal of the model itself is to minimize the mean squared error between the actual and predicted EEG responses. Model performance was assessed by calculating the correlation between the actual EEG of the held-out fold and the predicted EEG from that same fold using Pearson’s correlation coefficient (**Figure 1**). For the mismatch condition, we predicted EEG based on the stimuli they heard rather than the text they read.

Speech features can be highly correlated with one another, which makes it challenging to determine how each feature uniquely contributes to the neural responses to speech. In an effort to identify the unique contribution of each feature to those responses, we used a partial correlation approach, as we have done before (Synigal et al., 2023). Specifically, we fit and optimized five separate TRF models, one for each of the features mentioned above: envelope, spectrogram, phonetic features, lexical surprisal, and word onset. Then to identify the unique variance explained by any individual feature (Y), we used MATLAB’s *partialcorr* function (Fisher, 1924) to compute the partial Pearson correlation between the actual EEG data (X) and the predictions from the feature of interest’s model (Y), while controlling for the variance accounted for by the other four speech features (Z). This procedure was repeated for each feature model to quantify its unique neural contribution while accounting for the influence of the remaining features.

### Low-frequency event-related potential analysis

As well as modeling how the EEG tracked different speech features throughout the presentation of each specific stimulus, we were also interested to explore how the data might reflect more global differences in perception across trials. To that end, we also conducted an event-relate potential (ERP) analysis time locked to the onset of each degraded speech stimulus – with a focus on low-frequency activity that might reflect differences in perceptual evidence accumulation across trials. In particular, EEG epochs were extracted from the onset of each degraded speech sentence to the end of that sentence. Baseline correction was applied to each epoch by subtracting the mean of the EEG signal in the 500ms interval prior to audio stimulus onset. Before averaging across epochs, the epochs were truncated to a common length corresponding to the minimum duration across trials within each subject (2.05 ms). Artifact rejection was performed separately for each channel using a procedure based on peak-to-peak amplitude: for each epoch, the peak-to-peak voltage on a given channel was computed and converted to a median absolute deviation (MAD)-normalized z-score. For that channel, any trials exceeding a threshold of ±4 MAD-z were not included in the average. Remaining trials were categorized according to either experimental condition (Match vs. Mismatch) or behavioral rating (1–5) (see results). Within each subject, condition-specific averages were computed, as well as averages pooled across the two conditions for each rating level. These subject-level waveforms were subsequently used for quantification of ERP differences – again with a particular focus on low-frequency ERP dynamics.

### Statistical analysis

All statistical analyses were performed in MATLAB R2024b (MathWorks, 2024). One-tailed t-tests were performed on the behavioral scores. Two-tailed t-tests were performed on the partial correlation coefficients to assess deviations from zero and differences between conditions.

Permutation testing was performed to test the significance of the TRF model performance. A null distribution was generated by circularly shifting the stimulus features. Specifically, stimulus features were concatenated across trials, and a single random circular shift was applied to the concatenated time series. The same shift was applied to all stimulus features to preserve their temporal relationships. The minimum shift was set to exceed the maximum TRF lag (650 ms) to break the original stimulus–response alignment, while the maximum shift was set to be smaller than the total concatenated signal length minus the minimum shift to avoid near-complete wrap-around. Shifted stimuli were then segmented back into their original trial structure, and the TRF model for each feature was cross validated, trained, and tested. This procedure was repeated 1,000 times for both conditions.

Cluster-based permutation tests were then used to assess whether TRF prediction accuracies were reliably greater than chance across electrodes. For each subject and electrode, prediction accuracy was first centered relative to chance by subtracting the mean prediction accuracy obtained from the null distribution. Statistical inference was then performed on these chance-corrected prediction accuracies using within-subject, non-parametric cluster-based permutation tests implemented in FieldTrip. Clusters were formed by grouping neighboring electrodes that exceeded a non-parametric cluster-forming threshold (α = 0.05). Cluster-level statistics were computed by summing t-values within each cluster, and significance was assessed using 5,000 permutations. Monte Carlo methods were used to estimate p-values, with family-wise error rate controlled at α = 0.05 (Maris & Oostenveld, 2007). This procedure was repeated for each condition and feature.

For the ERP data, statistical analyses focused on two time intervals: 0.4 – 1.0 s and 0.4 – 2.0 s. Both intervals began at 400 ms when the sustained divergence between conditions observed in the grand-average waveforms began (please see results below). The first interval ended at 1.0 s to focus on the initial divergence between conditions and the other interval ended at 2.0 s to examine differences across the entire duration of the degraded speech (truncated to the shortest trials, as mentioned above). The mean and peak amplitudes of the ERP were computed across each of these time windows for each subject and condition and served as our primary dependent measures. Peak latency was also extracted within the narrower window, defined as time point at which the ERP reached its maximum in that window. In addition, a difference wave (ERP for match trials minus ERP for mismatch trials) was computed for each subject, and its mean amplitude within the analysis window was tested against zero. Group-level statistical comparisons between match and mismatch conditions were performed using paired-samples t-tests, and effect sizes were quantified using Cohen’s d for within-subject contrasts.

To assess the relationship between neural responses and subjective intelligibility, condition-averaged waveforms were further stratified by behavioral ratings (1–5). For each subject, mean amplitude within the analysis window was computed separately for each rating level. A linear trend across ratings was quantified by fitting a first-order polynomial to these values, yielding a per-subject slope estimate. These slopes were tested against zero at the group level using one-sample t-tests. In addition, a repeated-measures analysis of variance (ANOVA) was conducted across the five rating levels to assess overall modulation of the CPP by subjective report. All statistical tests were two-tailed, and significance was assessed at α = 0.05.

## RESULTS

### More words are heard and understood when degraded speech is preceded by matching text

In our task, participants listened to degraded speech and rated the number of words they were able to hear and understand on a scale of 1 to 5; 1 meaning they did not detect any words and 5 meaning they understood every word of a complete sentence. On average, participants reported hearing and understanding more words when the text prior matched the degraded speech content compared to the mismatch condition (t(27) = -28.229, p = 2.461 x 10^-21^, one-tailed t-test, **Figure 2**). The size of this effect was sizable (Cohen’s *d* = 5.395) and confirms that participants experienced a strong perceptual pop out during the match condition, consistent with previous research (Sohoglu et al., 2014).

**Figure 2.**
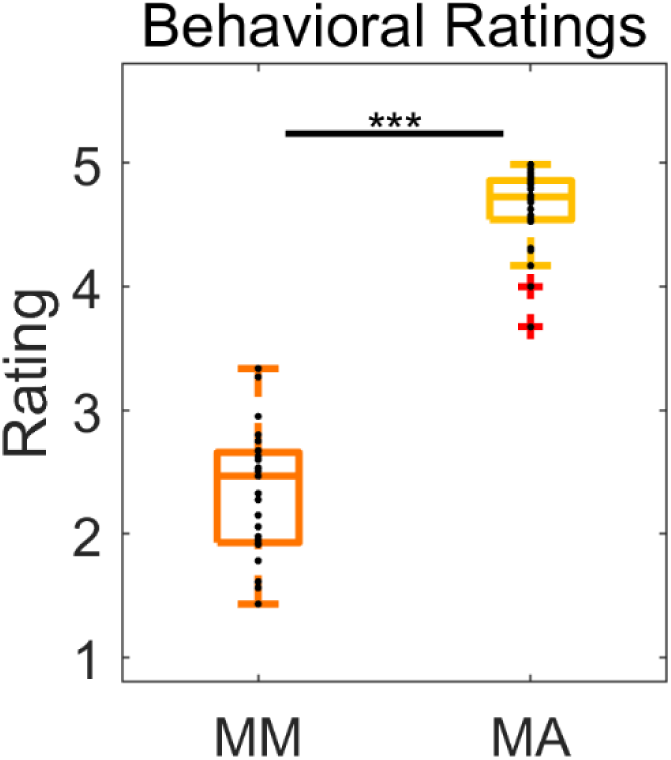
Behavioral results. Ratings of how many words participants were able to hear and understand in degraded speech clips that were preceded by matching or mismatching text. Each black marker represents a different participant. Significance is indicated by * if p < 0.05, ** if p < 0.01, and *** if p < 0.001.

### Phonetic feature processing may be influenced by prior information

The first goal of this study was to investigate how prior information alters acoustic and linguistic representations in the brain. Since participants were shown text primes before hearing the audio clips, we expected them to use this information to form predictions about upcoming speech. We hypothesized that acoustic tracking would not differ between the match and mismatch conditions, as the text did not provide a very precise prediction of how each audio clip would sound. In contrast, we expected differences in phonetic feature tracking, with lower prediction accuracies in the match condition. This expectation is derived from the predictive coding theory which proposes that a stimulus that matches a precise prior expectation leads to a smaller error signal and, thus, a weaker neural response. In our data, such a weaker response would lead to poorer prediction accuracy in our TRF model. In line with recent research (Karunathilake et al., 2023; Synigal et al., 2026), we also expected a difference in the performance of the lexical surprisal and word onset models between conditions. Specifically, because speech clarity is so much higher in the match condition, we expected to see evidence of neural responses time locked to word onset in that condition, but not the mismatch condition.

To assess how well the EEG tracked different aspects of the speech signal, separate TRF models were fit to the speech envelope, spectrogram, phonetic features, lexical surprisal, and word onsets. While similar information is encoded in the speech envelope and spectrogram, both were included to determine if neural tracking would reflect an average of all available information or a differential weighting of specific frequency bands. Model performance was quantified using prediction accuracy and visualized as scalp topographies (**Figure 3**). All feature-based TRF models significantly predicted EEG activity above chance in both the match and mismatch conditions, with significant positive clusters observed across all electrodes (all cluster-level p = 2.000 × 10^-4^, non-parametric cluster-based permutations). This pattern indicates robust neural tracking of each speech feature regardless of prior information type. In general, prediction accuracy was greatest over frontocentral scalp, which, given that our data were referenced to the average of the two mastoid channels, is consistent with activation of auditory cortex.

**Figure 3.**
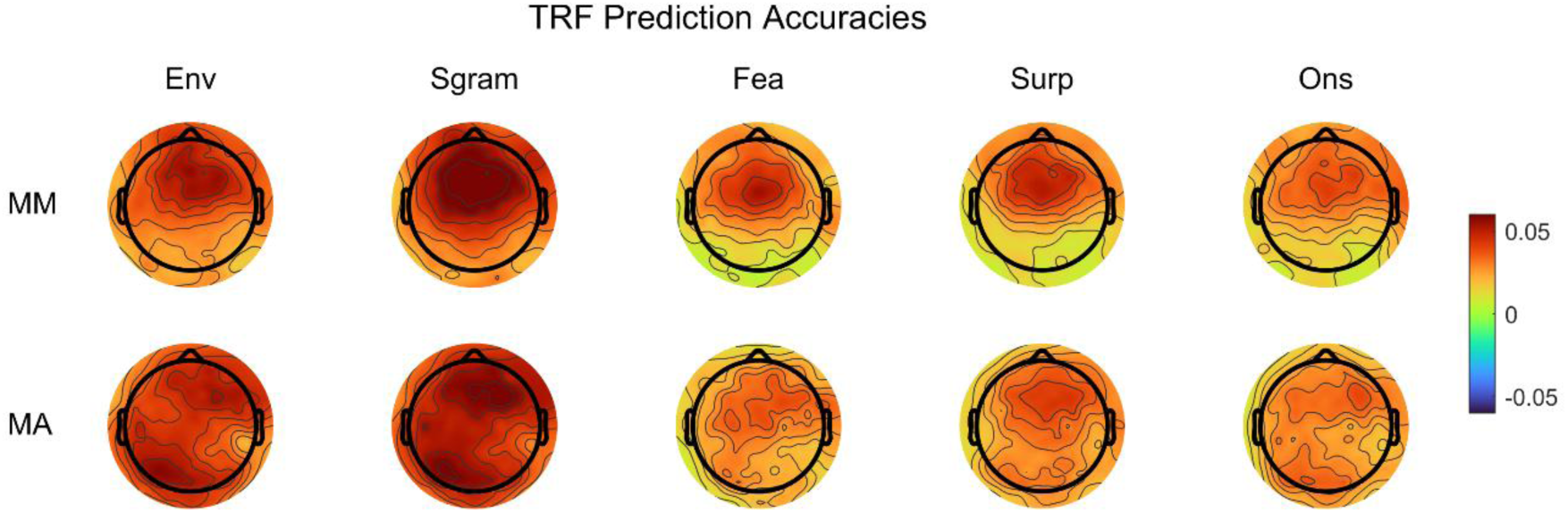
TRF model prediction accuracies. Scalp topographies show prediction accuracies for models based on the speech envelope (env), spectrogram (sgram), phonetic features (fea), lexical surprisal (surp), and word onsets (ons) at each electrode. Prediction accuracies reflect correlations between recorded EEG and EEG predicted by independently trained TRF models using each feature. Results are shown for the mismatch (MM; top row) and match (MA; bottom row) conditions.

The speech features we included in this study are correlated with one another and, as such, likely explain overlapping variance in the EEG data. Therefore, we computed partial correlation coefficients to assess the unique contribution of each feature. These coefficients were derived using the actual EEG and the predicted EEG obtained from independent TRF models. Specifically, we used MATLAB’s *partialcorr* function to compute the correlation between the actual EEG and the predicted EEG from one feature’s model, while controlling for the predicted EEG from all other feature models (**Figure 4**). One-tailed t-tests were used to determine whether the partial correlation coefficients were greater than zero for both conditions. In the mismatch condition, the spectrogram (t = 2.569, p = 0.008), phonetic feature (t = 5.050, p = 1.474 x 10^-5^), and lexical surprisal (t = 2.760, p = 0.005) coefficients were significantly greater than zero, whereas envelope (t = 1.065, p = 0.148) and word onset (t = 0.252, p = 0.402) coefficients were not. In the match condition, spectrogram (t = 3.591, p = 6.724 x 10^-4^) and lexical surprisal (t = 1.939, p = 0.032) coefficients were significantly greater than zero, while envelope (t = 0.096, p = 0.462), phonetic feature (t = 1.541, p = 0.068), and word onset (t = 1.189, p = 0.123) coefficients were not. These results indicate that the EEG reliably tracks the speech spectrogram and lexical surprisal, while phonetic features contribute more selectively in the mismatch condition. The fact that the envelope explained no significant variance in the EEG data makes sense given that whatever variance it might have explained is already accounted for by the spectrogram model. Similarly, the lack of any significant partial correlations for the word-onset model is not surprising because responses associated with word onsets are likely explained by the lexical surprisal model, while activity related to acoustic changes at word onset is likely explained by other features (e.g., the spectrogram).

**Figure 4.**
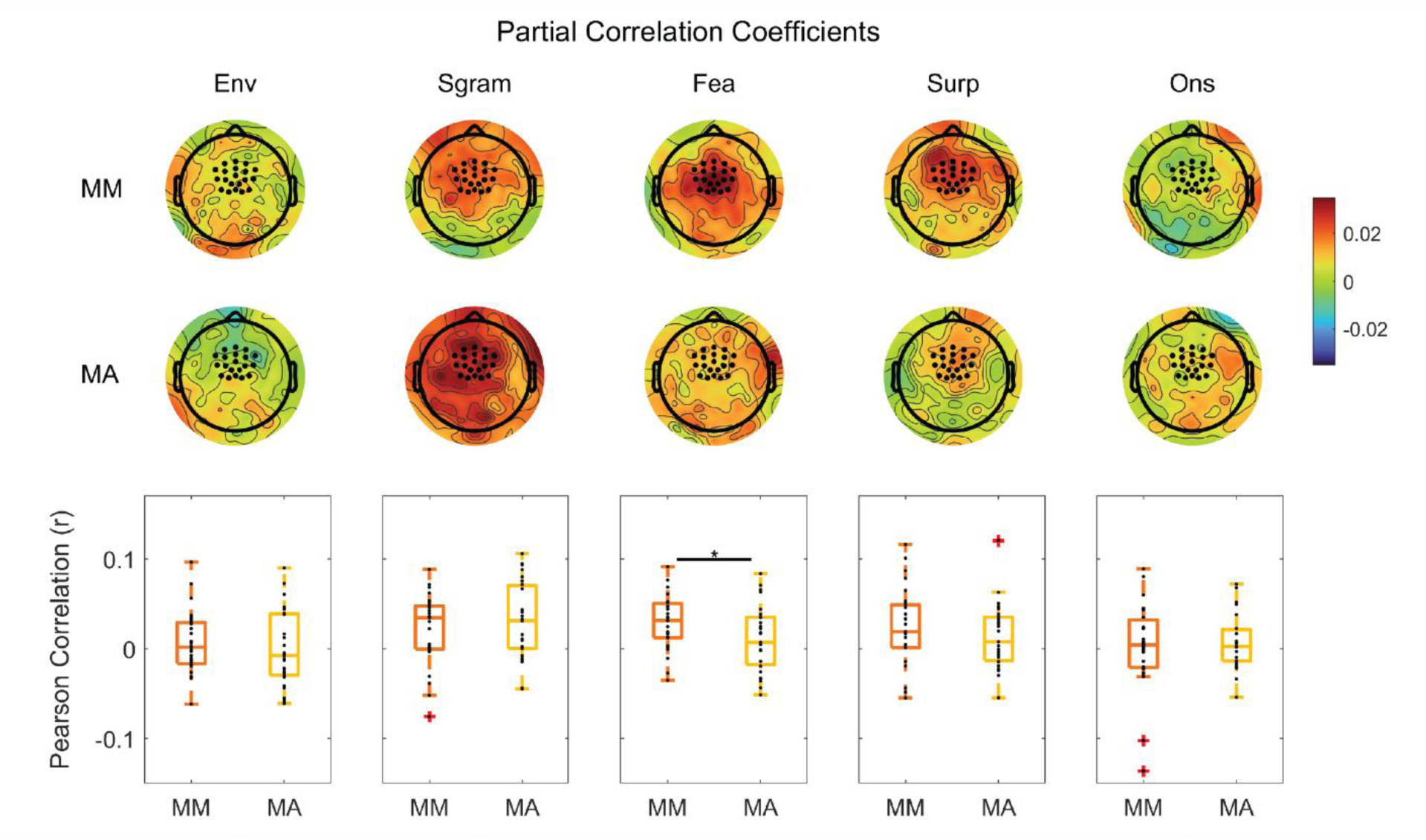
Unique contributions of each speech feature to the EEG responses. Scalp topographies show partial correlation coefficients for the speech envelope (env), spectrogram (sgram), phonetic features (fea), lexical surprisal (surp), and word onsets (ons) features at each electrode. Partial correlations reflect the unique contribution of each feature to the EEG, computed by calculating the correlation between the recorded EEG and the EEG predicted by that feature’s TRF model while controlling for predictions from all other feature models. Results are shown for the mismatch (MM; top row) and match (MA; bottom row) conditions. Boxplots show partial correlation coefficients averaged across 12 frontocentral electrodes (shown as black dots in the topographies) for each feature and condition. Each box represents the distribution across participants, with scores for individual participants being overlaid as black dots.

As expected, we found no differences in the envelope (t = 0.602, p = 0.552, two-tailed t-test) or spectrogram (t = -0.890, p = 0.382) partial correlation coefficients. More surprisingly, there were also no differences between the lexical surprisal (t = 1.045, p = 0.306) and word onset (t = -0.476, p = 0.638) partial correlations between conditions, despite the words being perceived much more strongly in the match condition compared to the mismatch condition. On the other hand, in line with our expectation, the phonetic feature model prediction partial correlation coefficients were significantly lower for the match condition than the mismatch condition (t = 2.267, p = 0.032) (**Figure 4**). Together, these results show that prior information selectively modulated phonetic feature tracking in our data, with reduced phonetic feature contributions when auditory input matches expectations.

While the effect on phonetic feature tracking is notable, three caveats are worth highlighting. First, the effect was very sensitive to preprocessing choices: the results reported above were obtained from EEG data detrended at 1 Hz but not high pass filtered. Applying a 1 Hz high-pass filter eliminated the observed differences between conditions (**Figure S1**), suggesting that very low-frequency signal components contribute to this effect and that that effect was not especially robust. Second, the reported effect reached significance (p = 0.032) without any correction for multiple comparisons. While our text-based prior led us to expect effects at the level of categorical phonetic feature processing and not at the level of acoustics (e.g., spectrogram) gives us some confidence that the effect is real, it is also true that it does not survive correction for multiple (partial correlation) comparisons. Finally, the size of the phonetic features effect reported above (**Figure 4**) was relatively small, with a Cohen’s *d* = 0.436. This was substaintially smaller than our behavioral effect size of 5.395.

To summarize our TRF model results: we find no evidence of an effect of (text) priors on the EEG tracking of the envelope, the spectrogram, lexical surprisal, or word onsets. We find some evidence of significantly weaker tracking of phonetic features for the match condition. However, this effect is not robust to different preprocessing choices, does not survive multiple comparisons tests, and displays a small effect size. All of this points to the idea that the encoding of the low-level acoustic and phonetic features of degraded speech are not strongly affected by prior information and, certainly, that any effects on such encoding are not of a similar magnitude to the very large effect size of the behavioral pop-out phenonemenon (**Figure 2**).

### A robust low-frequency neural signature of the effect of prior information on degraded speech perception

The lack of any large effects on any of our speech tracking measures above motivated us to search for neural indices that might more closely reflect the result of an inferential perceptual process. To that end, we examined low-frequency activity in ERPs time-locked to the onset of the degraded speech. This analysis revealed a striking difference between the match and mismatch conditions. Specifically, the ERPs in both the conditions displayed an auditory evoked potential to the onset of the degraded speech that did not differ between conditions, but this was immediately followed by a sharp divergence between conditions in the low-frequency EEG (**Figure 5**). This difference took the form of a positive increase in the EEG data for the match condition that began at about 400 ms (**Figure 5A**) and that was most prominent over midline parietal scalp (**Figure 5B**). Moreover, the difference in the average EEG traces between the two conditions persisted throughout the entire duration of the analysis window (up to 2 seconds), although it dropped a little after a peak at around 700 ms. Within the 0.4–1.0 s window, the mean amplitude was substantially higher in the match condition (2.2906 ± 3.1364) than in the mismatch condition (−1.5229 ± 2.8138). This effect was statistically significant across participants (*t*(26) = 8.2096, *p* = 1.085 x 10^-8^) and showed a very large effect size (Cohen’s *d* = 1.5799). Across the larger 0.4–2.0 s window, the mean amplitude was still substantially higher in the match condition (1.58 ± 3.46) than in the mismatch condition (−1.87 ± 3.63). This effect was also statistically significant across participants (*t*(26) = 4.92, *p* = 0.00042) and also showed a large effect size (Cohen’s d = 0.95). Peak amplitude analyses revealed a similar pattern, with larger peaks in the match condition (5.61 ± 4.08) relative to the mismatch condition (2.22 ± 4.01), which was significant (*t*(26) = 3.94, *p* = 0.00055) and showed a medium effect size (*d*= 0.77). For the sake of completeness, we also tested for any differences in peak latency between our conditions – although we had no particular predictions in this regard – and we found no difference between conditions in the 0.4-1.0s time window (match: 0.6983 ± 0.1600 s; mismatch: 0.7072 ± 0.2253 s; *t*(26) = -0.1940, *p* = 0.8477).

**Figure 5.**
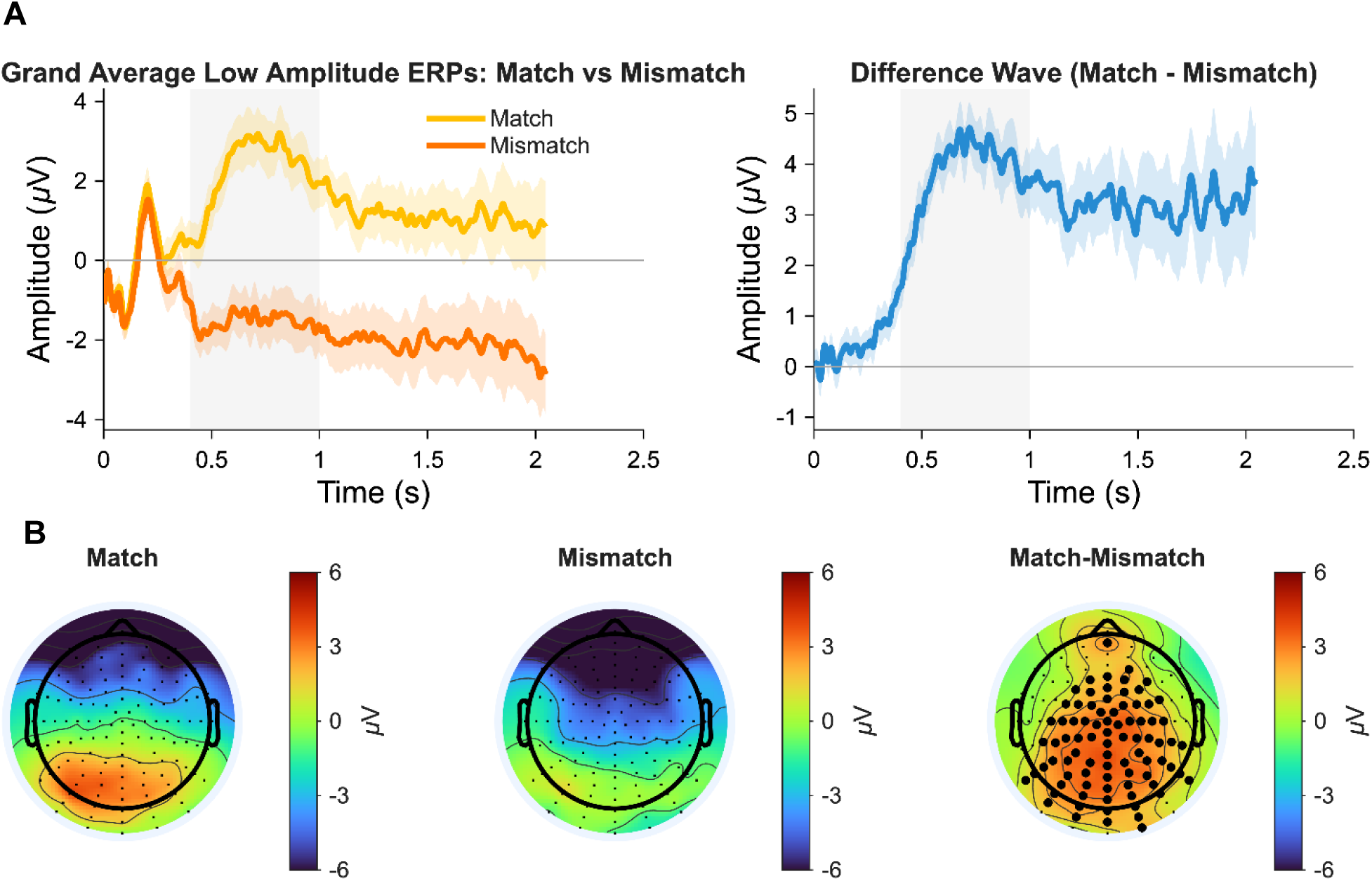
Group-averaged event-related potentials. (**A**) Grand-average ERPs for Match and Mismatch trials, showing an auditory evoked potential to the onset of the speech stimuli, followed by a steeper buildup for Match trials during the presentation of the degraded speech (left panel). Difference wave (Match − Mismatch), highlighting the steeply rising and prolonged positive difference between Match and Mismatch trials. Shaded region indicates the analysis window (0.4–1.0 s) used for computing the mean amplitude. (**B**) Topographic scalp distribution of grand-average event related potentials averaged across post-stimulus window 0.4–1 s when the two traces diverge. Topographies are shown for Match trials (left), Mismatch trials (middle), and their difference (Match − Mismatch; right), averaged across subjects. Electrodes marked with large black circles indicate channels showing a significant condition difference following false discovery rate (FDR) correction (q < 0.05).

### Low-frequency event-related potential activity correlates with perceived clarity

The above ERP analysis focused on comparing responses between our two main conditions – matched and mismatched priors. The results of that analysis are relatively interpretable given the sharp dichotomy in average behavioral responses between conditions (**Figure 2**). However, within and between participants there was also some variability in behavioral responses to individual trials. To more directly examine how the EEG data tracked perception at the level of trials, we computed grand average ERPs as a function of behavioral rating – pooled across both conditions (**Figure 6**). Visual inspection of the ERPs showed the steepest positive slope and most positive sustained low frequency activity for the trials that were rated most clear (a score of “5”), with trials rated “4” and “3” both appearing more positive than trials rated “2” and “1”. This pattern was confirmed by statistical testing that showed a significant positive linear relationship between behavioral rating (1–5) and larger ERP amplitude in the time window (0.4-1 s) (mean positive slope = 1.0508 ± 0.9515), *t*(26) = 5.7387, *p* = 4.8437 x 10^-6^, *d* = 1.1044. A significant positive relationship was also found for the 0.4 – 2.0 s window (mean positive slope = 0.97± 1.18), *t*(26) = 4.25, *p* = 0.000240, *d* = 0.82; not shown). Together, these results indicate that low-frequency event-related potential activity – centered over centroparietal scalp – is strongly modulated by prior congruency and scales with subjective perceptual clarity.

**Figure 6.**
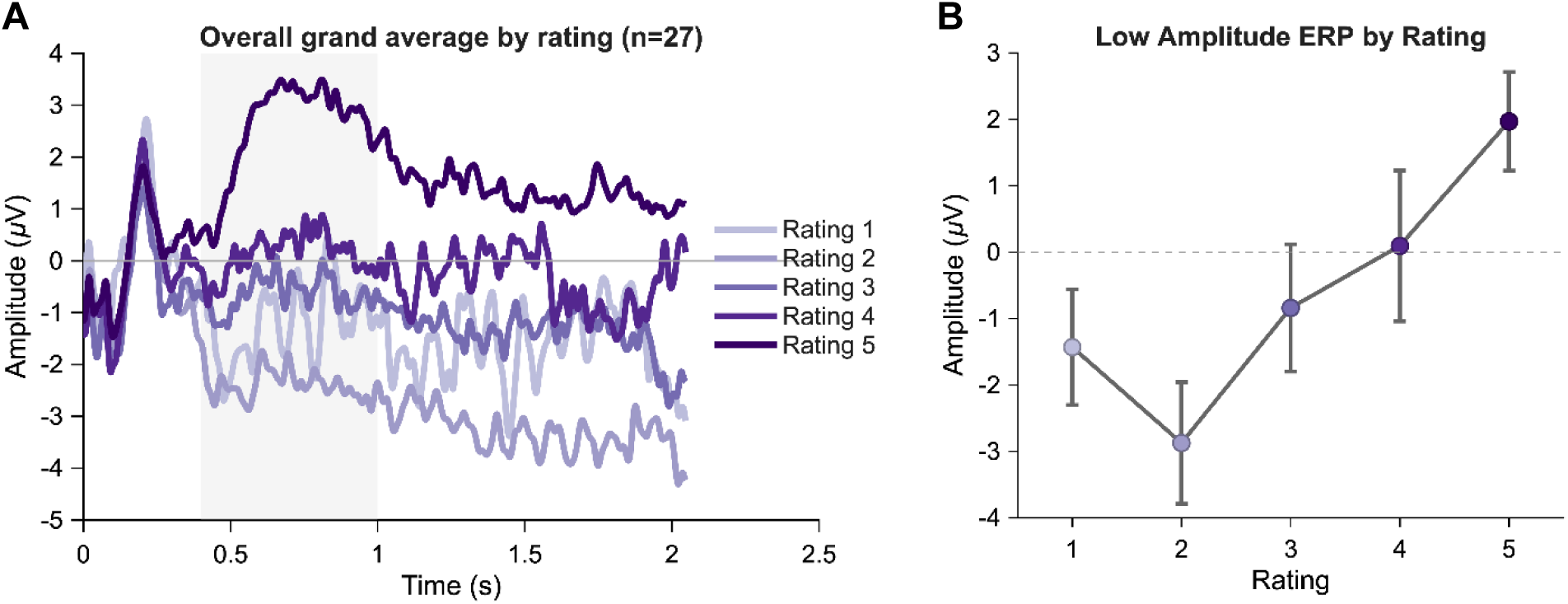
(**A**) Group-averaged ERP waveforms at electrode Pz as a function of subjective intelligibility rating (1–5), pooled across conditions. Shaded region indicates the analysis window (0.4–1.0 s). (**B**) Relationship between subjective intelligibility rating and ERP amplitude. Mean ERP amplitude at electrode Pz (averaged over the 0.4-1 s window) is plotted as a function of rating (1-5), showing an overall increase in amplitude with higher ratings. Error bars denote ±SEM across subjects. Trials perceived as more intelligible are associated with larger low-frequency ERP activity over midline centro-parietal scalp.

## DISCUSSION

This study investigated how prior knowledge influences the perception of degraded speech and how that influence is reflected in neural signatures of speech and language processing and perceptual inference. Participants were supplied with text priors that either were (match) or were not (mismatch) informative about the content of the upcoming degraded speech. In line with previous work (Sohoglu et al., 2014), we found a very large effect of prior information on perception, with degraded speech being perceived as much clearer when preceded by matching text. This was accompanied by a subtle effect on speech encoding in the form of weaker cortical tracking of phonetic features in the matched prior condition. It was also accompanied by a large effect in the form of a positive increase in the low-frequency EEG data over midline parietal scalp for the match condition that begins around 400 ms after stimulus onset and sustains throughout the stimulus. This positivity also scales with perceived clarity across trials.

A key goal of much perceptual neuroscience research is to identify interpretable patterns of neural activity that closely track with perception. In the context of the present study, we first sought to do this by examining how time-locked EEG measures of the acoustic and linguistic encoding of speech might reflect the very strong perceptual effect that comes with providing prior information ahead of the presentation of degraded speech. We reasoned that, because participants really feel that they are hearing the words in the match condition and not the mismatch condition, we might see striking between-condition effects on the cortical tracking of specific speech features. This turned out not to be the case. An initial analysis showed that the EEG tracked several individual speech features in both conditions (**Figure 3**). However, as these features are highly mutually correlated (Daube et al., 2019), interpreting the results of these tracking measures is not trivial, and, indeed, they are all likely to be dominated by general auditory processing (Prinsloo & Lalor, 2022). To deal with this issue, we also used a partial correlation approach to identify the unique variance in the EEG tracking measures that related to each of our selected speech features (**Figure 4**). The expectation here was that isolated signatures of low-level acoustic processing might look quite similar between match and mismatch conditions, given that both involve presenting speech with similarly degraded acoustics. However, we expected that isolated measures of speech-specific processing (e.g., the responses to phonetic feature categories and/or the responses to word onsets or word surprisal) might show large between-condition differences given the behavioral effect. The only effect we found seemed to be weaker tracking of phonetic features during the match condition, consistent with the possibility that such tracking reflects prediction error (Clark, 2013; Di Liberto, Crosse, et al., 2018; Friston & Kiebel, 2009; Sohoglu & Davis, 2020). That said, this effect did not survive multiple comparisons correction and was even sensitive to specific preprocessing choices, meaning that it falls short of being a pattern of neural activity that closely tracks with perception.

Our EEG speech tracking results add to a complicated picture as to how neurophysiological indices of degraded speech encoding are affected by prior information. For example, consistent with some previous studies, we find no evidence of stronger envelope tracking for our match condition (Di Liberto, Crosse, et al., 2018; Karunathilake et al., 2023), but counter to others (Baltzell et al., 2017; Corcoran et al., 2023). Meanwhile, our tentative result on phonetic feature encoding seems to be somewhat consistent with a previous study from our own group that showed a general reduction in phoneme encoding with prior knowledge (Di Liberto, Crosse, et al., 2018), but it was not seen in other work (Karunathilake et al., 2023). One way to potentially explain these inconsistencies is by considering a series of studies by Sohoglu, Davis, and colleagues (Sohoglu et al., 2024; Sohoglu & Davis, 2016, 2020). These studies involved manipulations of both the validity of prior information *and* of speech clarity and have consistently shown an interaction effect between validity and clarity. Specifically, they have shown that for more degraded speech, informative priors lead to larger neural responses, while for clearer speech, informative priors lead to smaller neural measures, a pattern that is consistent with those responses reflecting prediction error (Friston & Kiebel, 2009). As such, studies—including ours and many of those mentioned above—that do not manipulate speech clarity may be operating at a single speech clarity point where neural measures of speech processing could be larger, smaller, or no different as a function of prior validity. That said, some of the above studies did manipulate clarity and did not see an interaction effect (Baltzell et al., 2017). So even invoking the interaction of validity and clarity as a potential explanation for the inconsistencies in the above literature is somewhat tenuous.

Differences in other experimental factors between studies could also potentially explain the variability of findings based on cortical speech tracking. For example, some studies provide prior information in the form of clean audio (Baltzell et al., 2017; Di Liberto, Crosse, et al., 2018; Karunathilake et al., 2023) whereas others—including the present study—provide it in the form of text (Corcoran et al., 2023; Sohoglu et al., 2024; Sohoglu & Davis, 2020; Sohoglu et al., 2012). This necessarily entails a difference in the precision of prior information across hierarchical levels—with both providing precise predictions as to the upcoming phonemes, syllables, and words, but only the former providing very precise predictions as to the sounds. Another difference between studies centers on how often the stimulus is repeated: some studies presented degraded speech, followed by clean speech, followed by a repeat presentation of the degraded speech (Di Liberto, Crosse, et al., 2018; Karunathilake et al., 2023), while others, including our own, presented the degraded speech only once (Baltzell et al., 2017; Sohoglu et al., 2014). Given the long history of research showing that stimulus repetition can affect neural responses (Grill-Spector et al., 2006), this difference might also partially explain inconsistent findings. Another factor that varied across studies was the duration of the degraded speech stimuli, with some studies— including the present one—using single words or short, degraded sentences (< 5 s; (Baltzell et al., 2017; Banellis et al., 2020; Sohoglu et al., 2014)), and others using longer degraded speech segments (≥ 10s; (Corcoran et al., 2023; Di Liberto, Crosse, et al., 2018; Karunathilake et al., 2023)). This means that studies differ in terms of the demands placed on working memory for the prior information that might affect the precision of that prior information and its impact on perception. Moreover, it seems plausible that listening to degraded speech of different durations might also tax attention differently—something that is well known to strongly affect cortical speech tracking (Kerlin et al., 2010; O’Sullivan et al., 2015; Teoh et al., 2022; Zion-Golumbic et al., 2013).

While the question of how prior information influences the encoding of low-level acoustic-phonetic speech features is complicated, strong neural signatures of the perceptual pop-out phenomenon have been seen based on word level responses. For example, studies that have examined brain activity based on word onsets and/or word surprisal have reported clear evidence of a significant enhancement of that activity when prior information is available (Karunathilake et al., 2023; Synigal et al., 2026). Surprisingly, we did not find such an enhancement for either word onsets or word surprisal in the present study. The reason for this is not clear. However, previous work has shown that the cortical tracking of linguistic speech features can be attenuated when stimuli are brief, degraded, or provide limited sentential context for prediction (Brodbeck et al., 2018; Weissbart et al., 2020; Slaats et al., 2024). So, it is possible that the much shorter degraded speech stimuli used in the present study compared to those mentioned above might be the reason for our lack of an effect on word level responses. Indeed, using shorter stimuli might also have the effect of reducing (or eliminating) any responses based on word surprisal simply because the listener is not at all surprised by any of the words given that they had a perfect prediction of those words. The longer stimuli used in other studies may have been imperfectly remembered by the listener who might then have generated responses based on word surprisal during the degraded speech presentation.

As discussed above, the behavioral pop-out phenomenon is likely the result of a process of perceptual inference. Meanwhile, neurophysiological indices based on low-level features of the degraded speech stimuli are likely to relate much more closely to the initial sensory encoding of those features. The (often larger) word level response effects discussed above—because they cannot be explained solely based on the stimulus—are likely to more directly reflect perception. In the present study, we were also interested in identifying other neural signatures that might more directly index perception. With that in mind, we cast the problem as one of perceptual evidence accumulation and looked for signatures of such accumulation in the low-frequency EEG responses to the degraded speech stimuli. This analysis showed a marked difference between the conditions. This took the form of a positive increase in the EEG over midline parietal scalp beginning around 400 ms after stimulus onset and sustaining throughout the stimulus only for the match condition (**Figure 5**). This pattern is strikingly reminiscent of the centro-parietal positivity (CPP) (O’Connell et al., 2012) – an event-related potential measure that has often been reported as a signature of the accumulation of sensory evidence during perceptual decision-making (O’Connell et al., 2012; O’Connell & Kelly, 2021; Twomey et al., 2015). Precisely what such a CPP represents in our paradigm is not immediately clear. In typical perceptual decision-making tasks participants are asked to respond when they decide about the content of a sensory stimulus (e.g., that a pattern of dots is moving to the left or right). In our paradigm we did not specifically ask participants to make such a response, rather we asked them to listen to the entire stimulus and to rate its clarity afterwards. Given that our paradigm involved only two types of trial (match and mismatch), it is likely that participants recognized this early in the experiment. Thereafter, it is possible the participants settled on a strategy of determining whether the degraded speech stimulus did or did not match the preceding text. This would transform our clarity judgement task into a more binary decision-making task. If that were the case, however, one might wonder why there is no sign of a CPP during the mismatch trials when, presumably, participants would be accumulating sensory evidence towards deciding that the degraded speech does not match the text. The reason for this is not clear, although some recent decision-making work has suggested that CPP measures are significantly stronger when decisions are based on the presence of a task-relevant stimulus feature rather than the absence of that feature ({McCone, 2025 #1137}). The idea that our participants are making a decision based on whether the degraded speech matches or does not match the text would also be consistent with the fact that the CPP-like pattern in our data seems to scale with the reported clarity of the stimuli (**Figure 6**). When participants only hear a few of the words in the stimulus (e.g., clarity level 4 or 3), they have less perceptual evidence with which to make the decision about whether or not the stimulus matches the text. Across these stimuli then, the process of computing an ERP will likely mean averaging strong CPP responses to heard words with weak responses to words that were not heard leading to a lower amplitude response overall (**Figure 6**).

Another potential explanation for our pattern of results is that the low-frequency positivity in our EEG responses reflects the percept directly, rather than any specific decision based on that percept. Indeed, there is a long history of work using threshold stimuli that suggests that conscious perception involves “ignition” of the so-called global neuronal workspace, a process that is often characterized by the presence of a centro-parietal positivity in the form of the classic P300 response (Mashour et al., 2020). Given the very close relationship between the P300 (specifically, the P3b) and the CPP (O’Connell et al., 2012; Twomey et al., 2015), it may be that our data are directly reflective of ignition underlying conscious perception. Indeed, this idea has previously been advanced in the context of a pop-out paradigm involving monosyllabic words (Banellis et al., 2020), although that study reported a pattern that was more like a P3a response than the CPP-linked P3b. In any case, we suggest that it is a task for future work to disambiguate between direct neural markers of perception and those that are reflective of decisions based on perception. Such future work could include building a variety of tasks (including different decision-making tasks) around a perceptual pop-out paradigms like the one described in the present study.

Finally, we contend that the present findings have important implications for clinical research. For example, direct measures of conscious perception would be very valuable in assessing cognitive function in people with disorders of consciousness. This has previously been done using classic P300 paradigms (e.g., (Schnakers et al., 2009)). And it has also been done for word responses using both the classic N400 approach (Steppacher et al., 2013) and, more recently, continuous speech paradigms (Alkhoury et al., 2025). Identifying neural signatures of both word processing and perception using a pop-out paradigm could provide interesting complementary insights into the cognitive abilities of people with a disorder of consciousness. We also contend that our findings have implications for future research on psychosis. In particular, in recent years it has been hypothesized that hallucinations in psychosis might be the result of an overly strong weighting of prior information – relative to bottom-up sensory evidence – during perceptual inference (Cassidy et al., 2018; Corlett et al., 2019; Horga & Abi-Dargham, 2019; Kafadar et al., 2022; Powers et al., 2017; Teufel et al., 2015). Being able to index the neurophysiology of both sensory encoding and perceptual inference – and how those are affected by variations in the strength/validity of prior information – has great potential for testing this hypothesis. Indeed, the fact that most hallucinations come in the form of voices suggests that a speech-based paradigm – like the one used in the present study – might be a particularly effective approach.

## ACKNOWLEDGEMENTS

This work was supported by grants from the National Institute on Deafness and Other Communication Disorders (DC021140), the National Institute of Mental Health (MH135314), and the Del Monte Institute for Neuroscience at the University of Rochester. The authors thank Ms. Mariah Marrero for some assistance with the phoneme alignment.

